# On the accuracy of molecular simulation-based predictions of k_off_ values: a Metadynamics study

**DOI:** 10.1101/2020.03.30.015396

**Authors:** Riccardo Capelli, Wenping Lyu, Viacheslav Bolnykh, Simone Meloni, Jógvan Magnus Haugaard Olsen, Ursula Rothlisberger, Michele Parrinello, Paolo Carloni

## Abstract

Molecular simulations have made great progresses in predicting *k*_off_ values—the kinetic constant of drug unbinding, a key parameter for modern pharmacology—yet computed values under- or over-estimate experimental data in a system- and/or technique-dependent way. In an effort at gaining insights on this issue, here we used an established method to calculate *k*_off_ values—frequency-adaptive metadynamics with force field— and a subsequent QM/MM descriptions of the interactions. First, using force field-based metadynamics, we calculate *k*_off_ of the Positron Emission Tomography (PET) ligand iperoxo targeting the human muscarinic acetylcholine receptor M_2_. In line with previously performed *in silico* studies, the prediction (3.7 ± 0.7 ⋅ 10^−4^ s^−1^) turned out to differ significantly from the experimentally measured value (1.0 ± 0.2 ⋅ 10^−2^ s^−1^). Next, we use DFT-based QM/MM simulations to show that this discrepancy arises from erroneous force field energetics at the transition state. It turns out that this discrepancy is partly caused by lack of electronic polarization and/or charge transfer in commonly employed force field. We expect these issues to arise also in other systems where charged portions of the system play a pivotal role, such as protein- or DNA-protein complexes.

**Graphical TOC Entry:** 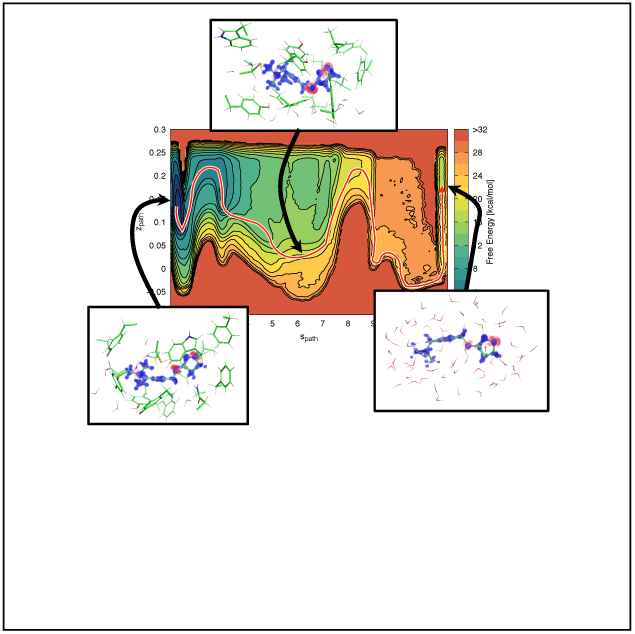

## Introduction

Efficacy and safety of drugs depends critically on their residence time. ^1,2^ Indeed, *k*_off_ values— the drug unbinding rate constant, corresponding to the inverse of the residence time— correlates with clinical efficiency even more than binding affinity. ^3,4^ Hence, the *k*_off_ value is one of the crucial parameters that current drug design strives to improve.^5,6^ While experiments face challenges to identify and characterize rate-limiting transition state(s), simulation approaches are able to predict free energy landscapes and residence times.^7^ Techniques devoted to this aim range from long-time molecular dynamics (MD) with specialized hardware,^8^ to a variety of different enhanced sampling methods such as random acceleration MD (RAMD),^9^ hyperdynamics, ^10^ conformational flooding, ^11^ Markov state models (MSMs)^12^ and infrequent^13^ or frequency adaptive metadynamics^14^ (I-MetaD, FA-MetaD). The latter two approaches have also predicted *k*_off_ values for ligands binding to cytoplasmatic proteins. All of the techniques mentioned above suffer from a systematic error of one to two orders of magnitude in comparison to the experiments. ^12,15–17^ This large discrepancy remains irrespective of the force field used (either Amber99SB^18^/GAFF^19^ or CHARMM22^*^ ^20^). This could be due to a variety of factors, including force field accuracy, molecular modeling procedures, and sampling issues. Here, we use a multistep simulation approach to address this important issue. We focus on the ligand iperoxo (Fig. 1 and SI1) that targets the human muscarinic acetylcholine receptor 2 (M_2_, Fig. 1), a system that is routinely used in neuroimaging and extensively investigated by some of us in a recent work. ^21^ First, we calculate the *k*_off_ value of the ligand by using the Amber14SB force field^22^ using well-tempered^23^ (WT) and FA-MetaD. To minimize errors due to the modeling procedure, we use the same pH and ionic strength as in the experimental conditions. ^24^ Once again, the predicted *k*_off_=3.7 ± 0.7 ⋅ 10^−4^ s^−1^ turns out to be much smaller than the experimental one (1 ± 0.2 ⋅ 10^−2^ s^−1^). Next, we compare the energetics of the force-field-based simulations with those based on combined quantum mechanics/molecular mechanics (QM/MM MD). The region treated at the quantum level consists of the drug and its interacting residues and it is described by density functional theory (DFT). The remaining part is treated as in the metadynamics calculation with the force field. There is a remarkable agreement between the *ab initio* and the force field estimation of the ligand/protein binding energy (Fig. 1). However, this is not the case for the transition state. Our analysis indicates that the lack of polarization is one of the key factors causing this discrepancy, which in turn affects the accuracy of the k_off_ calculation.

**Figure 1:**
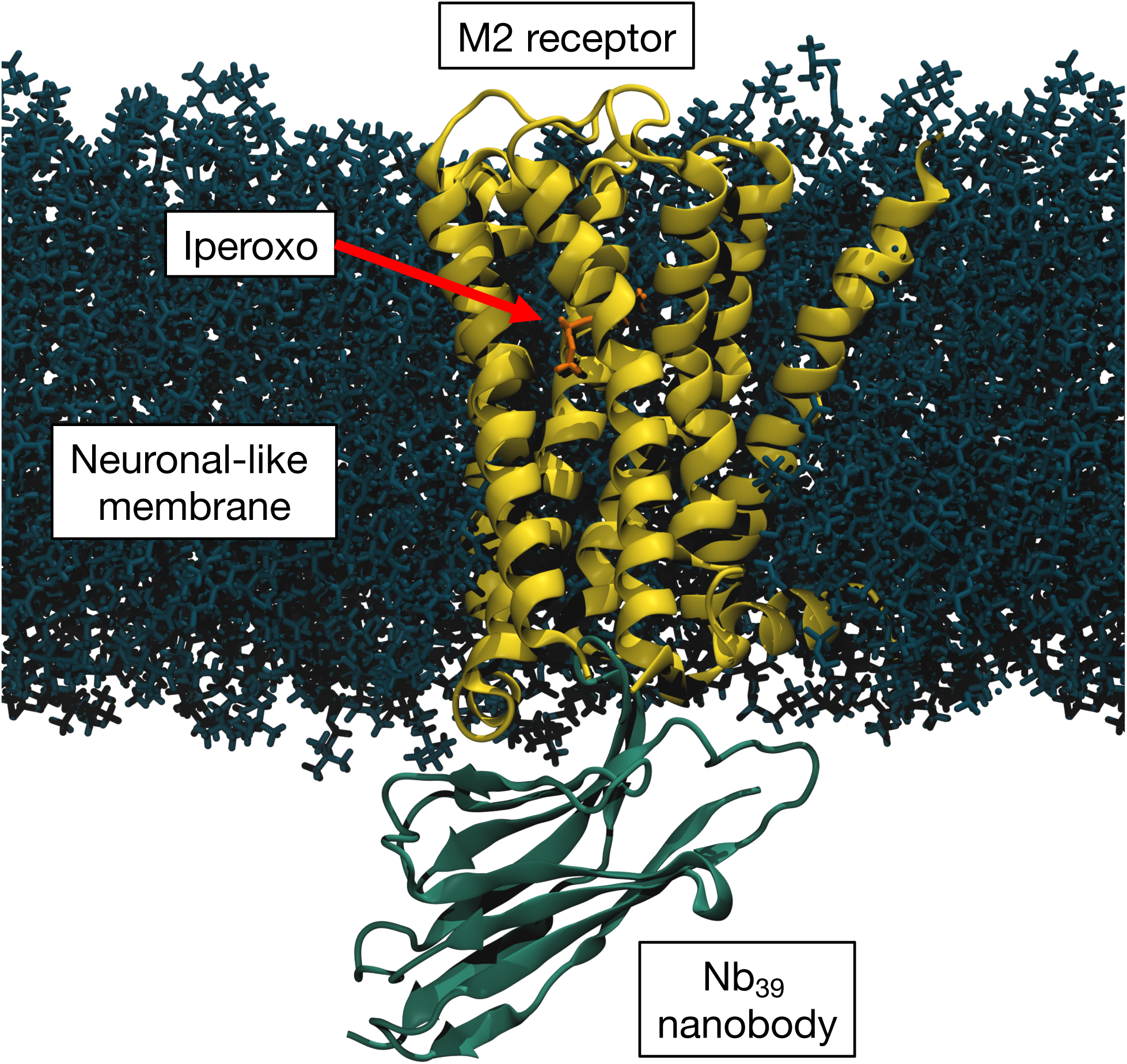
Representation of the simulated system. The M_2_ receptor (yellow), with its agonist iperoxo (orange), is embedded in a neuronal-like membrane^26^ (dark blue) and it is bound to the Nb_39_ nanobody (green) which keeps it in its active state. Water and Na^+^/Cl^−^ solvated ions are removed for sake of clarity.

## Results

Recently, WT funnel^25^ MetaD, based on the classic AMBER14SB force field ^22^ (CFF hereafter) predicted the full free energy landscape of the ligand iperoxo binding/unbinding pathways in the M_2_ muscarinic receptor in its active state^21^ (SI, Fig. 2). This prediction is a prerequisite for MetaD-based calculation of k_off_. Two different possible binding/unbinding pathways emerged (here noted with **I** and **II**). In pathway **I**, the ligand starts from the bound state, rotates around the axis formed by the alkyne bond, passing through the transition state 1 (TS1, Fig. 2), to finally reach state **A**. After this step, we observe a rotation of the entire ligand with the trimethylammonium group as a pivot (transition state 2 - TS2), ending in state **C**. A slightly different rotation around the same pivot can lead to state **B**. This second rotation does not lead to unbinding, and will not be investigated here. After reaching state **C**, the trimethylammonium group breaks the salt bridge formed with D103 (TS3) and moves toward the extracellular part, reaching the fully solvated state. The rate-limiting step is TS2. Pathway **II** is identical to **I** until the ligand reaches state **C**. Here, the rearrangement of the extracellular loop 2 (ECL2) of the receptor limits the possibility of the ligand to reach the solvated state right after visiting state **C**, forcing it to perform a further rotation, reaching the last metastable state **D**. After that rotation, the ligand breaks the salt bridge formed with D103 and it moves towards the solvent, completing its pathway to solvation.

**Figure 2:**
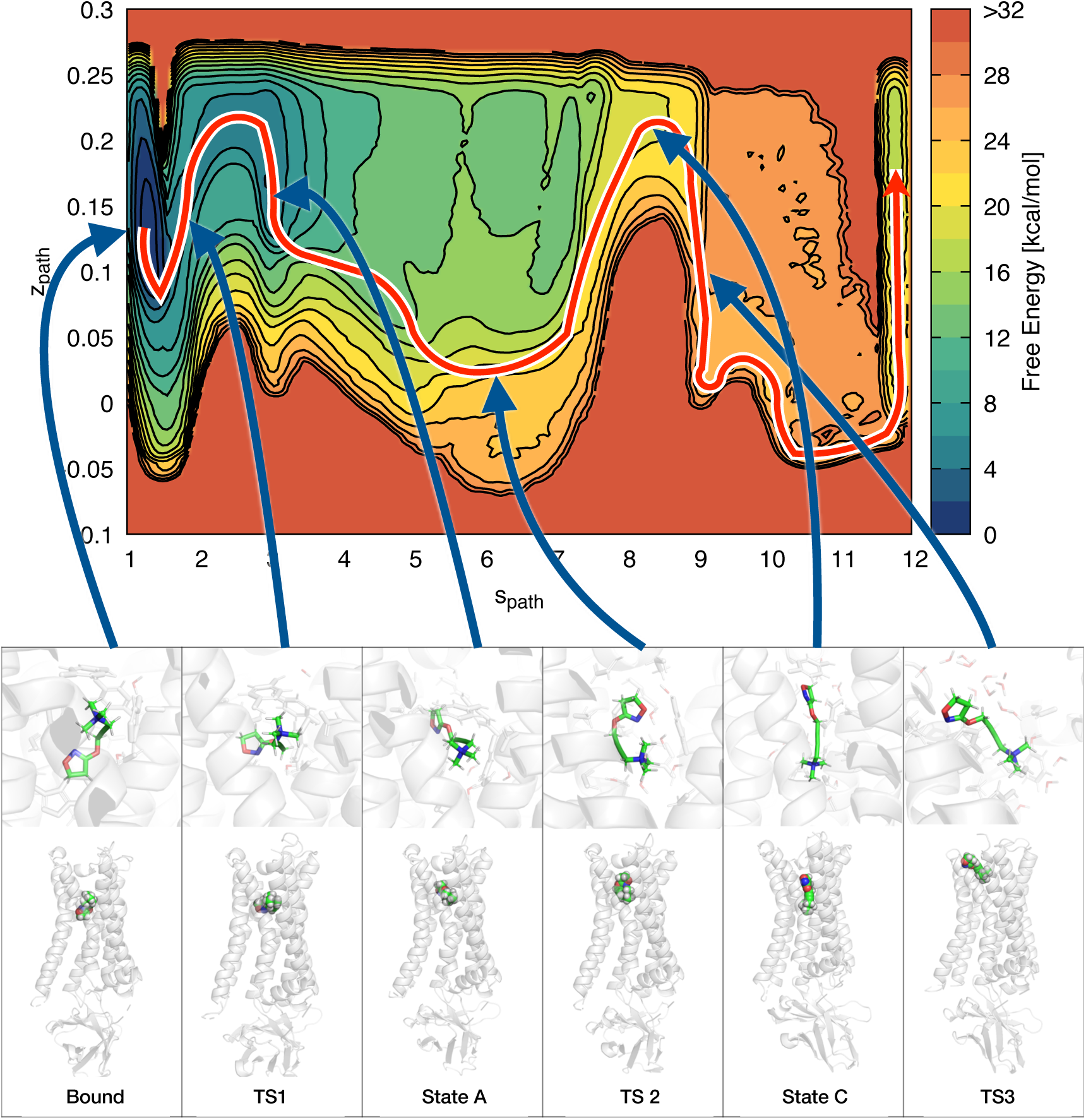
Free energy surface of binding with the observed unbinding pathway and representative structures. The top part shows the M_2_/iperoxo free energy surface as a function of *s*_path_ and *z*_path_, with the unbinding pathway followed by the ligand. The bottom part represents the Bound, TS1, A, TS2, C, and TS3 states: in the upper panel, both the ligand and its surrounding atoms within 4.5 Å are rendered; in the lower panel, the ligand is rendered in sphere mode with all the receptor and the nanobody. Water and ions are not represented for sake of clarity.

Here we evaluate the *k*_off_ by a multistep approach as in Casasnovas *et al.*^15^ Because kinetic calculations are very expensive, we explore only pathway **I** using appropriate path collective variables (pathCVs).^27^ This particular kind of CV is meant to study a single pathway between two reference states, limiting the motion of the system only around this predefined path. In the pathCV approach, two variables *s*_path_ and *z*_path_ are defined, where *s*_path_ defines the progression along the pathway, while *z*_path_ samples deviation from the reference path (in our case **I**). Next, we perform WT-MetaD ^23^ to calculate the free energy as a function of *s*_path_ and *z*_path_. Finally, we use FA-MetaD, ^14^ for the actual calculation of *k*_off_.

### Choice of pathCVs

The definition of a pathCV is based on a metric that measures the distance of instantaneous configurations from the path. In the first applications of pathCV,^27^ the metric chosen was based on root mean square displacement (RMSD) with respect to the initial bound state. In our case, having seen the presence of multiple intermediate states along pathway **I**^21^ (Fig. 5), we prefer to define our metrics based on a contact map based on the ligand–protein atom pairs that are crucial for the stabilization of the intermediate states (see SI for detail). To obtain a sequence of conformations along the unbinding trajectory, we employed Ratchet&Pawl MD^28,29^ in a two step-approach. First, we forced our system to perform its unbinding transition using as CV the distance between the binding pocket and the center of mass of the ligand (in the same spirit as a previous work^30^); from these structures we built a first pathCV. As a second step, we apply Ratched&Pawl to this first pathCV. From the set of conformations obtained during this run, we built the final pathCV that we employed in the MetaD simulations.

### WT MetaD

Multiple-walker WT MetaD as a function of *s*_path_ and *z*_path_ was carried out with 10 replicas. After a total simulation time of 1.8 *µ*s, we observed a large number of recrossing events and we computed the free energy surface along pathway **I** (Fig. 2). Although the CV was slightly different, the unbinding process was the same as in our previous work^21^ (Fig. 2 in the SI).

### FA-MetaD

We performed 10 different FA-MetaD^14^ runs, biasing both *s*_path_ and *z*_path_. Out of 10 runs, 5 were removed because they deposited bias on the transition state, invalidating the sampling performed.^13^ The 5 production runs covered 0.9 to 1.7 *µ*s, for a total simulation time of ∼ 8*µ*s.

The distribution of calculated residence times obtained (Fig. 3) is fitted with a Poisson distribution. We obtained a residence time 2.7 ± 0.5 ⋅ 10^3^ s and a *k*_off_ = 3.7 ± 0.7 ⋅ 10^−4^ s^−1^. To validate the correctness of the calculation performed, we performed a Kolmogorov–Smirnov test between the obtained distribution and the theoretical one.^31^ The p-value turned out to be 0.87. This shows that the obtained distribution is statistically indistinguishable from a theoretical rare event distribution. Hence, the discrepancy compared to the experimental value (*k*_off_ = 1.0 ± 0.2 ⋅ 10^−2^ s^−1^)^24^ does not seem to result from limited sampling but must be ascribed mostly to the underlying potential energy function.

**Figure 3:**
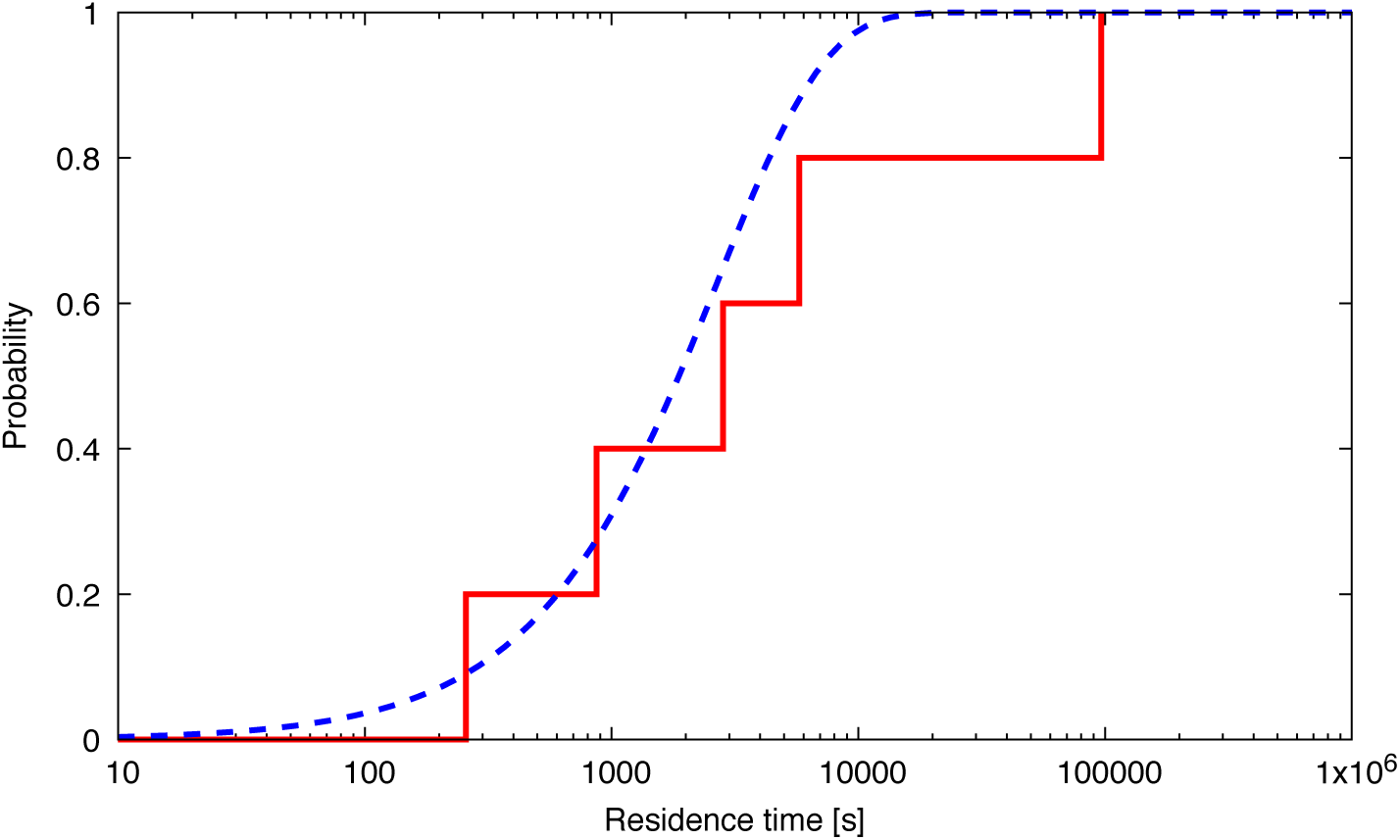
Comparison between the calculated (red line) and the theoretical Poisson distributions (blue dashed curve).

In order to validate the above hypothesis, the ligand/protein energetics is recalculated at the quantum mechanics level for selected configurations along the dissociation path. Specifically, we calculate the ligand/receptor interaction energies of the ligand at the bound and TS2 states considering the unbound state (the ligand in solution) as our reference state (i.e., computing the ∆∆*E* with respect to the solvated state). To make these computationally feasible we use a QM/MM approach as implemented in the MiMiC multiscale interface.^32,33^ The QM part of the system consists of the ligand and its interacting groups (Fig. 5a-c, See Methods for details). It is treated at the DFT level, using either the B3LYP^34–36^ or the BLYP^34,35^ exchange-correlation functionals. The rest of the system is described by CFF as in the metadynamics study. The energy of the total system is calculated by adding QM/MM interaction energy to the QM and MM energies, while that of the ligand is given by the QM energy. The ∆∆E values are also calculated at a purely MM level using the CFF for comparison.

In the bound state ∆∆E, calculated for the same 10 representative conformations, turn out to be very similar using the force field and DFT (−17.3±1.5 kcal/mol (CFF) and - 18.1±3.1 (B3LYP)/-18.1±3.1 (BLYP) kcal/mol), i.e. the values are not significantly affected by the exchange–correlation functional (Fig. 4, see also Table 2 in SI). By increasing the number of representative conformations to 45, the values for CFF and BLYP do not change (Fig. 4). Taken together, these results are consistent with previous WT-MetaD-based free energy calculations which showed excellent agreement between calculated and experimental affinities for this^21,30^ and other systems. This is expected and it is indeed confirmed by countless examples, both in protein–ligand^37^ and protein–protein^38^ interactions. This further supports the conclusion that modern force-field-based calculations can accurately predict ligand binding affinities.

**Figure 4:**
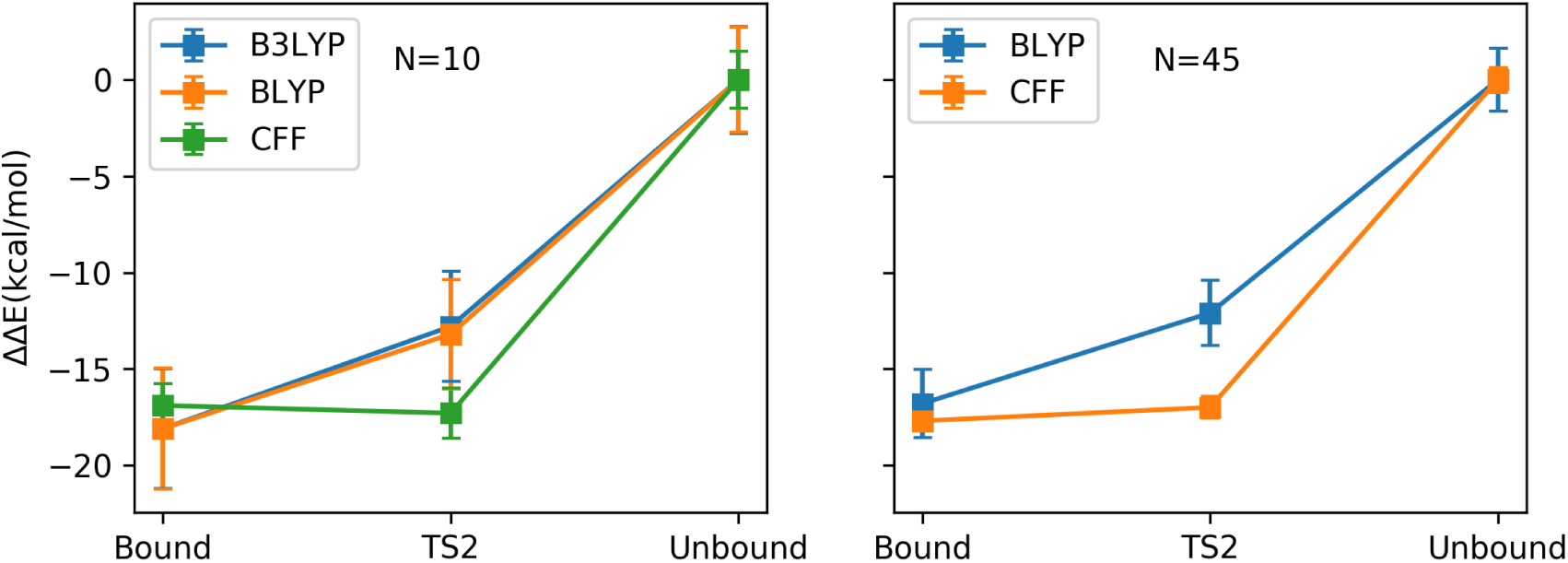
Relative interaction energies (∆∆*E*) of the ligand to the M_2_ receptor at QM/MM level (with BLYP and B3LYP exchange–correlation functionals) and CFF level. N is the number of conformations considered in statistics for each state. The ∆∆*E* of the interaction energies is not directly related to the enthalpy of the binding/unbinding process; in particular, a direct comparison between the different states cannot be performed for the different number of atoms considered.

A dramatically different scenario takes place at the transition state of the unbinding process, TS2. Here the value obtained with the force field (−17.8 ± 1.3 kcal/mol) differs significantly from the DFT ones, while the latter are still similar to each other (−12.8 ± 2.9 (B3LYP)/-13.2 ± 2.8 (BLYP) kcal/mol). The trend is preserved when the number of conformations is increased. Correcting the free energy barrier represented by TS2 with this enthalpic contribution would lead to a better representation of the energetics, and lower the residence time. However, also the entropic contribution to the free energy of the transition state may be affected by the limited accuracy of the potential energy surface of force fields. Thus, here the discussion must be kept at a qualitative level. Consistently with Haldar and coworkers, ^39^ present results suggest that simulations are suitable to compute unbinding kinetics provided that the dynamics is driven by sufficiently accurate forces.

Finally, we investigate if changes in electronic polarization ^40^ and charge transfer (CT) effects (which are lacking in routinely used biomolecular force fields) occur in the unbinding process. To achieve this, we calculate, using QM/MM, the rearrangement of electronic density of the ligand while passing from *in vacuo* to the bound state, to the transition state TS2, and to the unbound state (Fig. 5 a-c). These turn out to involve mostly the positively charged trimethylammonium group (forming a salt bridge with Asp103 or with water) and the two oxygen atoms (Fig. 5 a-c). As expected, the overall CT (∆*Q*_*CT*_) as well as the polarization (∆*Q*_*Pol*_), albeit small in magnitude (a fraction of an elementary charge), are significant for the three states (Fig. 5 d). Importantly, at the two stable states (bound and unbound), the ∆*Q*_*CT*_ shows a good consistency (both are 0.11 ± 0.01e); while it is about 30% larger on passing from the bound/unbound states to TS2 (0.14 ± 0.01e). Obviously, the fixed charge schemes of commonly used force fields cannot take into account these changes in charge redistribution during the unbinding processes.

**Figure 5:**
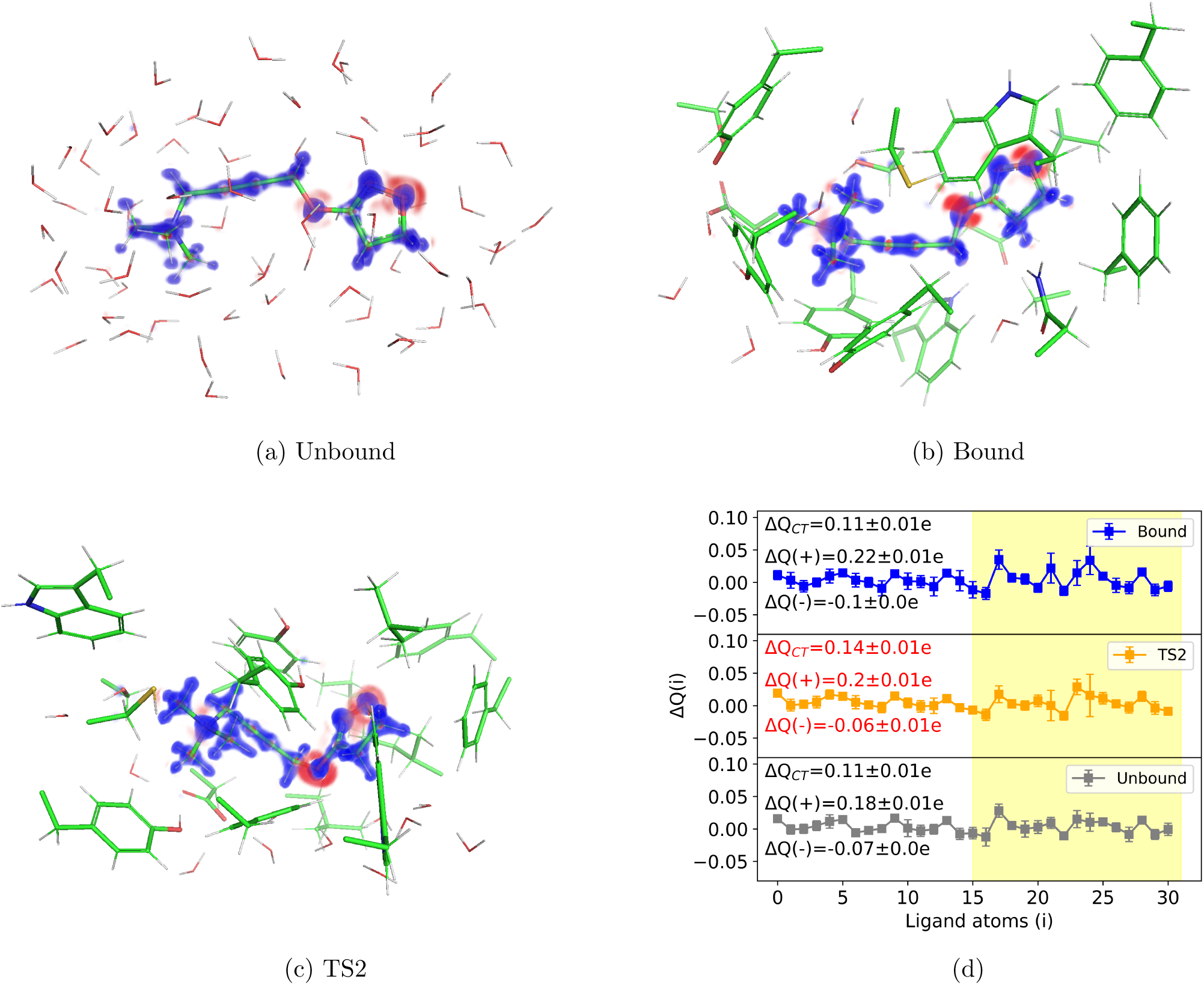
(a-c) Change of electronic density of the ligand on passing from *in vacuo* to in water (∆*ρ*= *ρ*^*cmplx*^ − *ρ*^*lig*^ − *ρ*^*rest*^) for the unbound (a), bound (b), and TS2 (c) states (blue: ∆*ρ* > 0, red: ∆*ρ* < 0). The atoms are displayed in sticks mode. (d) Correspondent change in atomic charge for each atom (∆Q(i), i= 0-30). ∆Q_*CT*_ is the overall charge transferred from the protein to the ligand, ranging from 0.11 to 0.14 electrons. ∆Q(+) and ∆Q(−) are the polarizations of the ligand contributed from atoms with positive and negative ∆Q(i), respectively, ranging from −0.1 to 0.22 electrons. The yellow color band highlights the most significant differences among the three states.

## Conclusions

The *k*_off_ constant is exponentially related to the height of the free energy barrier for dissociation. Small errors in the force field may therefore introduce large errors in the *k*_off_ of drugs—a key parameter in pharmacology. Here, we take iperoxo bound to the M_2_ receptor, a real life system used clinically, and we find that, while the bound state is excellently described, the accuracy of the force field at the transition state is limited,^39^ at least in part, by the lack of electronic polarization and charge transfer effects. These effects are present and slightly change when the ligand passes from the bound to the transition state and cannot be captured by standard non-polarizable force fields. Polarizable force fields, ^41–43^ reactive MD,^44^ and/or corrections of the free energy landscape derived from quantum mechanical calculations ^39^ might alleviate this problem. Similar issues may be expected in other unbinding processes, such as those involving protein/protein and protein/DNA complexes.

## Methods

### System preparation

We followed the same protocol as in ref. ^21^ to build and equilibrate the systems and run MD. Briefly, we obtained the structure from the protein data bank (PDB code: 4MQS^45^), parametrized the ligand using GAFF,^19^ and embedded it in a neuronal-like^26^ membrane. Then the system was solvated, ions were added to reach the experimental ionic force, and finally minimizing and equilibrating the system (details in the SI). All the simulations were performed using GROMACS 2018.4.^46^ patched with PLUMED 2.5.^47^

### rMD Simulation and pathCV identification

To identify a collection of conformations to set up our pathCV variable, ^27^ we used Ratchet&Pawl MD.^28,29^ As an initial ratcheting coordinate, we considered the distance between the center of mass of our ligand and the center of mass of the pocket (defined by TYR104, SER107, VAL111, PHE195, and TYR239), projected along the axis normal to the membrane. We fixed the bias factor to *k* = 500 kJ/mol/nm and the final ratchet coordinate to *r*_final_ = 2.5 nm. After 10 different 20-ns long rMD runs with these parameters, we selected 11 frames that describe well the progression of the ligand toward the solvent, and we used them to define a pathCV based on the contact map^48^ between a list of atom pairs of the ligand and the receptor (list in the SI). With this variable, we performed 10 new 20-ns long rMD runs to verify and eventually refine the new variable, choosing again by visual inspection 11 different frames from the unbinding trajectories and redefining a final pathCV that was then used in our MetaD simulation.

### WT MetaD along the pathCV

Using the pathCV as identified above, we performed multiple-walkers^49^ WT MetaD^23^ along the *z*_path_ and *s*_path_ using 10 different walkers. We set the bias factor to 24, an initial Gaussian height of 1.2 kJ/mol, and a frequency deposition of 1 ps^−1^. To limit the phase space exploration to path **I** only, we prevented our system from reaching pathway **II**, by putting a restraint to avoid its motion along that pathway (i.e. we put a harmonic wall at *z*_path_ = 0.25, where paths **I** and **II** diverge). The total simulation time was 1.8 *µ*s. The free energy surface was reweighted *a posteriori* with the Tiwary and Parrinello algorithm. ^50^

### FA-MetaD

We carried out 10 different FA-MetaD^14^ runs. The approach is a variant of I-MetaD^13^ that speeds up the calculations (details in SI). We performed 10 different FA-MetaD runs, with a bias factor of 24, an initial Gaussian height of 1.2 kJ/mol, an initial frequency deposition of 1 ps^−1^, an acceleration parameter θ = 100 (details in the SI), and a minimum frequency deposition of 10^−2^ ps^−1^. Out of all simulation performed, 5 have deposited bias on the transition state, and thus we discarded them, obtaining the 5 residence times from the remaining simulations.

### QM/MM single point calculations

A selection of 45 structures associated with the Bound, TS2, and Unbound states (Fig. 2) underwent 1000 steps of energy minimization using the steepest descent algorithm at the CFF level. Then, for each structure, we considered the *total system*, the *rest* (i.e. the system without the ligand) and the *ligand* without the systems (i.e. in vacuum). Overall, 135 structures were considered.

The QM regions in the *total system* consisted of the ligand, and the side-chains (up to the -Cβ) directly interacting with it as well as water molecules within 4.5 Å from it. They ranged from 196 to 308 atoms (see Table 3 in SI). The QM regions of the *rest* were the same except that the ligand was not included. They ranged from 165 to 277 atoms. Those of the *ligand* included only the latter (31 atoms).

The QM part was described at the DFT level (QM part, Fig. 2), using either the B3LYP or BLYP exchange–correlation functional. ^51^ A plane-wave basis set with a cutoff of 90 Ry was used. The core electrons were described through norm-conserving Troullier–Martins pseudopotentials.^52^ Isolated system conditions i were achieved by using the Martyna–Tuckerman scheme.^53^ For the Bound and TS2 states, (i) covalent C_*α*_-C_*β*_ bonds across the QM/MM interface were described by an adapted monovalent carbon pseudopotential;^54^ (ii) the net charge of the residues considered at QM level were reweighted to their sidechain atoms by neutralizing the sum of the partial charges of the remaining backbone atoms in MM region. For the *total system* and *rest*, the atoms other than those in the QM regions (’MM region’) were described using exactly the same setup and force field as for the metadynamics. The interactions between the QM and MM parts were described as in ref. 32: (i) the electrostatic interactions were calculated explicitly for MM atoms that are within 30 Å of the centroid of the QM part, whereas the interactions with the rest of the system was evaluated using a 5th-order multipole expansion of the electrostatic potential; ^32^ (ii) the Grimme’s correction ^55^ was used to describe dispersion interactions.

Single point electronic structure calculations were performed, with a convergence criteria of 10^−7^ au, using the highly scalable MiMiC-based QM/MM interface,^32,33^ which combines CPMD 4.1^56^ with GROMACS 2019.4. ^57^

The ligand binding energy ∆*E* was calculated either at QM/MM (∆E(B3LYP/BLYP)) or at the force field (∆E(CFF)) level. It reads

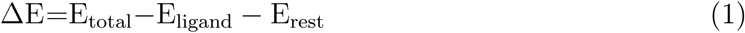

where *E*_total_ and *E*_rest_ are the potential energies of the *total* system and of the *rest* (given by summing the QM energy with the MM energy and the QM/MM interaction energies), and *E*_ligand_ the potential energy of the *ligand*.

We calculated ∆E at BLYP and CFF levels for all of the 45 conformations. Then, we chose 10 structures covering the same spreading range of the calculated energies at the BLYP level for the more expensive and accurate B3LYP calculations ^1^.

The change in electron density upon ligand binding was calculated at the the B3LYP level:

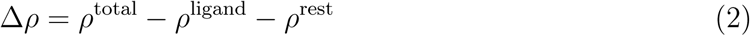

Here, *ρ*^total^ is the electron density of the QM part embedded in the MM part of the *total system*, *ρ*^ligand^ is that of the *ligand* and *ρ*^rest^ is that of the *rest*. The latter contribution exists only for TS2 and bound states.

The electron charge transfer (CT) associated with atom *i* of the ligand reads:^58^

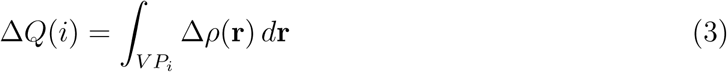

The integral is solved numerically over the grid points within the Voronoi partition of atom *i* (*V P*_*i*_) as in ref. 58. The CT effect of the whole ligand molecule reads

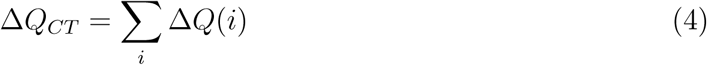

An estimation of the change in charge distribution is given by electric polarization as:

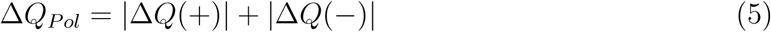

where, ∆*Q*(+)= Σ_*i*_ ∆*Q*(*i*), *i* ∈ {∆*Q*(*i*) > 0} and ∆*Q*(−)= Σ_*i*_ ∆*Q*(*i*), *i* ∈ {∆*Q*(*i*) < 0}.

## Acknowledgement

The authors thank Anna Bochicchio and Rodrigo Casasnovas for the initial preparation of the system, Emiliano Ippoliti, Luca Maggi, and GiovanniMaria Piccini for useful discussion. The authors gratefully acknowledge the computing time granted through VSR on the supercomputer JURECA^59^ at Forschungszentrum Jülich (Project ID: jias5d) and acknowledge the JSC for the computing time on the supercomputer JURECA Booster module. This project has received funding from the European Union’s Horizon 2020 Research and Innovation Programme under Grant Agreement No. 785907 (HBP SGA2) and the Marie Sklodowska-Curie Grant Agreement No. 642069. W.L. appreciates the National Natural Science Foundation of China No. 21505134. J.M.H.O. acknowledges financial support from the Research Council of Norway through its Centres of Excellence scheme (Project ID: 262695). U.R. acknowledges funding from the Swiss National Science Foundation via the NCCR MUST and individual grants.

## Supporting Information Available

All the data and PLUMED input files required to reproduce the results reported in this paper are available on PLUMED-NEST (www.plumed-nest.org), the public repository of the PLUMED consortium, ^60^ as plumID:20.005.

Details on Ratchet&Pawl MD, FA MetaD, system preparation and choice on the CV and on the QM/MM calculations are available in the Supporting Information.

## Footnotes

1 For comparison, the statistical estimate at BLYP and CFF were re-evaluated for the 10 structures.

