## Supplemental Information for "On the accuracy of molecular simulation-based predictions of k_off_ values: a Metadynamics study"

### Supplementary Information

#### Supplementary Table 1: Contact map atom pairs

List of the pairs that form the contact map used in the pathCV definition. If a list of atoms is inserted, we consider the center of mass of those. The atom names for the protein residues are the standard ones from the Amber14SB force field.[1] The names of ligand atoms are given in Supplementary Figure 2.

| ID | atoms | protein residue | iperoxo atoms |
|---|---|---|---|
| $d_1$ | CG,CD1,CE1,CZ,CE2,CD2 | TYR262 | N1 |
| $d_2$ | CG,CD1,CE1,CZ,CE2,CD2 | TYR239 | N1 |
| $d_3$ | CG,CD1,CE1,CZ,CE2,CD2 | TYR104 | N1 |
| $d_4$ | OD1,OD2 | ASP103 | N1 |
| $d_5$ | CG,CD1,NE1,CE2,CZ2,CH2,CZ3,CE3,CD2 | TRP236 | C8,N2,O2,C10,C9 |
| $d_6$ | CB,CG1,CG2 | VAL111 | C8,N2,O2,C10,C9 |
| $d_7$ | CG2,CG1,CD1 | ILE178 | C8,N2,O2,C10,C9 |
| $d_8$ | CB,SG | CYS176 | N1 |
| $d_9$ | CB,CG2,OG1 | THR187 | C8,N2,O2,C10,C9 |

### Supplementary Table 2: Ligand/protein interaction energies at the QM/MM and force field (FF) levels

The intermolecular interaction energies $\Delta E$ (equation 1) of the ligand in different states along the unbinding pathway are calculated at the QM/MM (B3LYP/BLYP) and CFF levels. The differences in interaction energy $\Delta\Delta E$ are defined with respect to the unbound state (i.e. $\Delta\Delta E = \Delta E$ - $\Delta E$(unbound) ). All the reported values are in kcal/mol.

| | N | Bound State | Transition State (TS2) | Unbound State |
|---|---|---|---|---|
| $\Delta E$(B3LYP) | 10 | -127.6 ± 2.4 | -122.3 ± 2.1 | -109.5 ± 2.0 |
| $\Delta E$(BLYP) | 10 | -124.7 ± 2.5 | -119.8 ± 2.1 | -106.6 ± 1.9 |
| $\Delta E$(CFF) | 10 | -52.5 ± 0.5 | -52.9 ± 0.8 | -35.6 ± 1.0 |
| $\Delta\Delta E$(B3LYP) | 10 | **-18.1 ± 3.1** | **-12.8 ± 2.9** | **0 ± 2.8** |
| $\Delta\Delta E$(BLYP) | 10 | **-18.1 ± 3.1** | **-13.2 ± 2.8** | **0 ± 2.7** |
| $\Delta\Delta E$(CFF) | 10 | **-17.3 ± 1.5** | **-17.8 ± 1.3** | **0 ± 1.5** |
| $\Delta E$(BLYP) | 45 | -123.2 ± 2.9 | -118.5 ± 2.6 | -106.4 ± 2.4 |
| $\Delta E$(CFF) | 45 | -52.7 ± 0.3 | -52.0 ± 0.3 | -35.0 ± 0.4 |
| $\Delta\Delta E$(BLYP) | 45 | **-16.8 ± 1.8** | **-12.1 ± 1.7** | **0 ± 1.6** |
| $\Delta\Delta E$(CFF) | 45 | **-17.7 ± 0.5** | **-17.0 ± 0.6** | **0 ± 0.6** |

### Supplementary Table 3: Details of the QM/MM systems.

Atoms treated at QM level in the bound, unbound and the TS2 states, respectively. This number changes depending on the selected MD snapshot. The average value, standard deviation, minimum value, and maximum value of the total number of QM atoms (Natoms), the number of QM atoms belong to protein (Nres), the number of QM atoms belong to water molecules (Nsol) are reported, respectively.

| | Natoms | Nres | Nsol |
|---|---|---|---|
| *Bound* | 248 ± 15 | 196 ± 13 | 21 ± 5 |
| *TS2* | 262 ± 16 | 205 ± 13 | 26 ± 7 |
| *Unbound* | 219 ± 9 | 0 ± 0 | 188 ± 9 |
| *Bound(min)* | 210 | 170 | 9 |
| *TS2(min)* | 235 | 186 | 18 |
| *Unbound(min)* | 196 | 0 | 165 |
| *Bound(max)* | 304 | 246 | 27 |
| *TS2(max)* | 308 | 228 | 39 |
| *Unbound(max)* | 238 | 0 | 207 |

### Supplementary Note 1: Ratchet&Pawl MD

Ratchet&Pawl Molecular Dynamics (rMD)[2,3] is an out-of-equilibrium sampling technique. Starting from one conformation of the system, we define a scalar ratcheting coordinate $r(t)$ that can drive the system toward a second state of interest (reaching the ratcheting coordinate $r_{\text{final}}$ that corresponds to the end point). We then apply a bias that dumps thermal fluctuations in the direction opposite to the end point. This is achieved by imposing the following "ratchet" potential:

$$V_{\text{rMD}}(\xi(t)) = \begin{cases} \frac{k}{2}\left(\xi(t) - \xi_m\right)^2 & \xi(t) > \xi_m(t) \\ 0 & \xi(t) \leq \xi_m(t) \end{cases} \tag{1}$$

where

$$\xi(t) = (r(t) - r_{\text{final}})^2$$

and

$$\xi_m(t) = \max_{t \in [0,T]} \xi(t)$$

### Supplementary Note 2: Infrequent and Frequency Adaptive Metadynamics

In the case of zero deposition of bias in the transition state, metadynamics allows to predict the timescale of a transition between two different states of the system.[4] To achieve this, one deposits bias with a slower rate than standard metadynamics (i.e. infrequent metadynamics), giving the possibility to the system to have a transition between those deposition. In these conditions, the simulated time $t_{\text{sim}}$ in a metadynamics run turns out to be related to the real time $t$ with the equation

$$t = \alpha t_{\text{sim}}, \tag{2}$$

where $\alpha$ is the *acceleration factor*, and can be computed from the bias deposited by metadynamics $V(s, t_{\text{sim}})$ as

$$\alpha = \langle \exp(\beta U(s, t_{\text{sim}}) \rangle \tag{3}$$

where $\beta = (k_B T)^{-1}$, $s$ is the position of the system in the CV space at time $t_{\text{sim}}$, and $< \cdot >$is the average over all the simulation.

After a certain number of different metadynamics runs, we obtain a set of residence times of the ligand inside the receptor. To verify that we are simulating a rare event and no bias is deposited on the transition state. (i) we fit on our empirical distribution the theoretical cumulative distribution function (CDF) of a homogeneous Poisson process of characteristic time

$$\text{CDF}_{\text{Poisson}} = 1 - \exp(-t/\tau) \tag{4}$$

to the empirical CDF of our residence times. (ii) We can perform a Kolmogorov-Smirnov test between the fitted rare event CDF and the residence times empirical CDF: if the p-value is >0.15, we can safely assume that the obtained times are extracted from a rare event distribution.[5]

Frequency Adaptive metadynamics[6] is a variant of Infrequent Metadynamics. There, the deposition frequency is not fixed, but it is adaptively slowed down when the acceleration factor $\alpha$ grows. This change permits to speed up the bias filling of the initial free energy minima, allowing to reduce the simulation time, and maintaining the "infrequent" behavior near the transition states. Practically, one choose an initial deposition $\tau_0$, an upper deposition time limit $\tau_c$ (that have to be in the same order of a usual infrequent metadynamics deposition time), and a parameter $\theta$, that drives the dynamics of the deposition time. The deposition time $\tau_{\text{dep}}$ at time $t$ reads

$$\tau_{\text{dep}} = \min\left(\tau_0 \cdot \max\left(\frac{\alpha(t)}{\theta}, 1\right), \tau_c\right) \tag{5}$$

where $\alpha(t)$ is the instantaneous acceleration factor, defined as

$$\alpha(t) = \exp(\beta U(s(t), t) \tag{6}$$

### Supplementary Note 3: Details on System Preparation

Our preparation protocol is the following. The X-ray structure of the $M_2$/iperoxo complex in presence of a nanobody was obtained from the Protein Data Bank (code: 4MQS[7]). The lipids composition was chosen to represent that of a model neuronal membrane[8] (i.e., 48% cholesterol, 16% phosphatidylcholine (DPPC and POPC), 16% phosphatidylethanolamine (DOPE), 14% sphyngomielin (SM 18:0), 4% phosphatidylserine (DOPS), and 2% phosphatidylinositol (SOPC)). The receptor was inserted into membrane bilayers following the orientation of the OPM database.[9] The system was then solvated with explicit water molecules. We neutralized the charge of the system and added a salt concentration of 150 nM, following the condition used in experiments.[7,10] The final simulation box was 112 x 113 x 149 Å$^3$, containing around 150.000 atoms in total. The parametrization of the model is the same as in our previous work,[11] except for the calculation of the atomic charges for iperoxo: before the calculation of the charges with the restrained electric potential fitting method (RESP)[12] at HF/6-31G* level of theory, we performed a further geometric optimization at B3LYP/6-31+G(d,p) level of theory to have a better estimation of the single-point charges employing Gaussian 09.[13] The simulations were performed using GROMACS 2018.4.[14] patched with PLUMED 2.5.[15] The protein, membrane, counterions, and water were described by the AMBER ff14SB force field,[1] the CHARMM-GUI[16] and Slipids,[17,18] Joung and Cheatham force fields,[19] and TIP3P,[20] respectively. We used the generalized AMBER force field (GAFF)[21] for the ligand. A cutoff distance of 12 Å was used for the van der Waals and short-range electrostatic interactions, and the long-range electrostatic interactions were computed with the particle-mesh Ewald (PME) summation method[22] using a grid point spacing of 1 Å. Long-range dispersion corrections to the pressure and potential energy were considered.[23]

Our equilibration protocol started with the lipid tails: restraining all the other atoms, the lipid tails were energy minimized for 1000 steps using the steepest descent algorithm and subsequently melted with an NVT run for 0.5 ns at 310 K. After this initial run, we activated pressure coupling for a 10 ns run, equilibrating the entire system to a stable volume, fixing only the positions of the protein and the ligand. In this equilibration phase, we noticed that after the first 5 ns the volume of the box reached a plateau, fluctuating around its equilibrium value without any canonical drift. We then released the protein restraints for a further 0.5 ns keeping the pressure of 1 bar and physiological temperature (i.e., 310 K). After these minimization and equilibration procedures, we performed a long unrestrained MD simulation for 0.7 $\mu$s. The temperature was kept constant using velocity rescaling thermostat[24] with solvent, solute, and membrane coupled to separate heat baths with coupling constants of 0.5 ps. The pressure was maintained constant with a semi-isotropic scheme, so that the pressure in the membrane plane was controlled separately from the pressure in the membrane normal direction, and the Parrinello-Rahman barostat[25,26] was applied with a reference pressure of 1 bar, a coupling constant of 2 ps, and a compressibility of $4.5 \cdot 10^{-5}$ bar$^{-1}$ for both the directions.

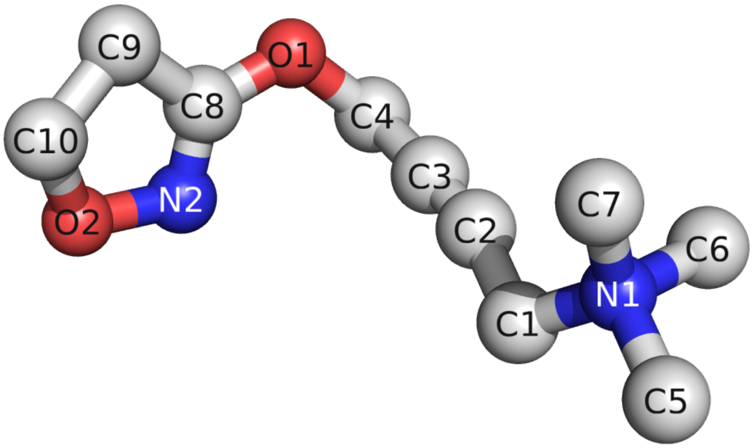


Supplementary Figure 1: Iperoxo molecule in ball and stick representation. On every atom we put the atom name used in our parameterization. The hydrogen atoms are not shown for sake of clarity.

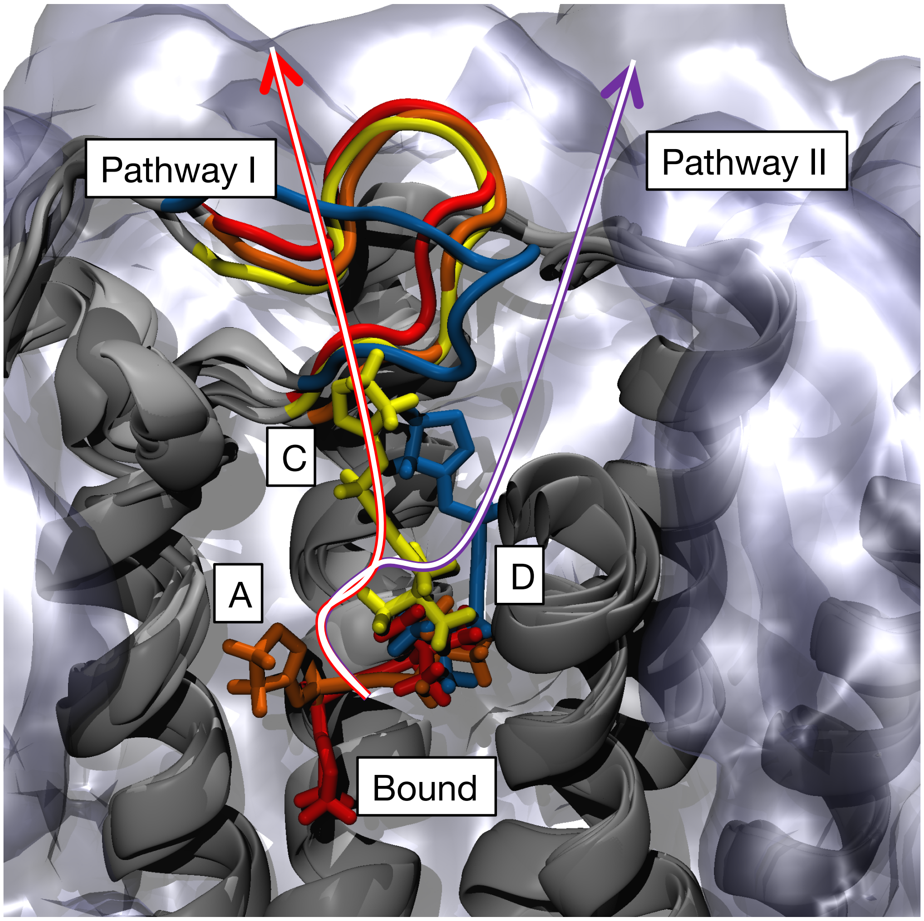


Supplementary Figure 2: Visual representation of the two unbinding pathways of iperoxo ligand from the $M_2$ receptor as identified in ref.[11] Iperoxo (stick representation) starts from the Bound state (red) and for both the pathways **I** (red arrow) and **II** (violet arrow) passes through state **A** (orange) and **C** (yellow). If the extracellular loop 2 (ECL2) is the crystallographic position (yellow, orange and red), the ligand can make a translation toward the solvent, reaching the fully solvated state (pathway **I**); if ECL2 is rotated, it hampers the motion of the ligand, that is forced to reach the intermediate state **D** (blue), moving then toward the solvent.
